# GEOMetaCuration: A web-based application for accurate manual curation of Gene Expression Omnibus metadata

**DOI:** 10.1101/257444

**Authors:** Zhao Li, Jin Li, Peng Yu

## Abstract

Metadata curation has become increasingly important for biological discovery and biomedical research because a large amount of heterogeneous biological data is currently freely available. To facilitate efficient metadata curation, we developed an easy-to-use web-based curation application, GEOMetaCuration, for curating the metadata of Gene Expression Omnibus datasets. It can eliminate mechanical operations that consume precious curation time and can help coordinate curation efforts among multiple curators. It improves the curation process by introducing various features that are critical to metadata curation, such as a back-end curation management system and a curator-friendly front-end. The application is based on a commonly used web development framework of Python/Django and is open-sourced under the GNU General Public License V3. GEOMetaCuration is expected to benefit the biocuration community and to contribute to computational generation of biological insights using large-scale biological data. An example use case can be found at the demo website: http://geometacuration.yubiolab.org. Source code URL: https://bitbucket.com/yubiolab/GEOMetaCuration

## Introduction

Metadata curation is an essential step for analyzing and integrating heterogeneous large-scale biological datasets generated by numerous labs across the world (1). Such data integration presents biomedical researchers with unprecedented opportunities to discover new biological insights that are hidden if each dataset is dealt with separately (2). However, it becomes increasingly challenging to curate semi-structured metadata efficiently and accurately, as the volume of biological data grows rapidly (3). For example, there are more than 100,000 datasets in one of the most popular public databases for functional genomics data, Gene Expression Omnibus (GEO) (4). It is critical to develop new methods to facilitate curation to ultimately enable streamlined biological discovery from a large amount of biological datasets.

Despite the importance of metadata curation in biological discovery, the topic has received relatively little attention compared with other areas of data curation. Only recently, using techniques from natural language processing (NLP), Bernstein *et al.* (5) developed a computational pipeline to automatically recognize biomedical entities of human biological samples from metadata in the National Center for Biotechnology Information’s (NCBI’s) Sequence Read Archive (SRA). However, curating high-level summary information is still an unsolved problem in NLP (6), such as determining the key factors being perturbed in a dataset. Manually curating these metadata is essential. Indeed, a number of metadata resources, such as RNASeqMetaDB (7) and SFMetaDB (8), have been created using manual curation with high accuracy.

However, manual metadata curation is time-consuming. General-purpose applications, such as Google Drive, have been used to facilitate the curation process to a certain extent (7) (8). But the curation process still involves many mechanical and tedious operations interspersed throughout. Curators are required to collect information scattered in multiple places for each curation task to reach a correct conclusion. Much time is spent searching in these places to locate relevant fragments of text. In addition, due to potential errors made by each curator, a curation task must be assigned to multiple curators, and their curation results must be compared and integrated to ensure high curation accuracy. Such a coordinated effort can take a significant amount of time without the assistance of a tool.

To address these problems with metadata curation, we developed a curator-friendly web-based application, GEOMetaCuration, for GEO datasets. It was designed to eliminate mechanical steps in the curation process to improve curation productivity for a large amount of metadata. The application facilitates curation in two ways: 1) a well-designed back-end management system simplifies the process of coordinating multiple curators and multiple curation topics; and 2) an integrated curation page for various dataset information and automatic keyword highlighting helps curators improve their efficiency. By improving the productivity of manual curation, GEOMetaCuration is expected to benefit the biocuration community and to contribute to computational generation of biological insights using large-scale biological data.

## System implementation

GEOMetaCuration is designed to facilitate the manual curation process of GEO metadata, including the following steps: 1) downloading from GEO the metadata of an initial list of datasets for a specific curation topic; 2) assigning the metadata of each dataset to multiple curators; 3) (curators) curating the metadata by examining relevant materials including dataset descriptions, sample descriptions, abstracts of corresponding papers, and sometimes full papers; and 4) collecting intermediate results from curators and finalizing them (**Figure 1**). Initially, the GEO dataset metadata for a curation topic are downloaded by E-Utilities (9) and are imported to a database in GEOMetaCuration. On the GEOMetaCuration administration webpage, the metadata are assigned to curators. To verify curation consistency, the metadata of each dataset are curated by multiple curators. Curators curate their assigned datasets independently using the web interface, and the results are stored in the database. GEOMetaCuration then automatically collects intermediate curation results for all datasets. Results are cross-checked for integration, and only consistent results are recorded. Inconsistency in the remaining intermediate results are resolved by the administrator.

**Figure 1.**
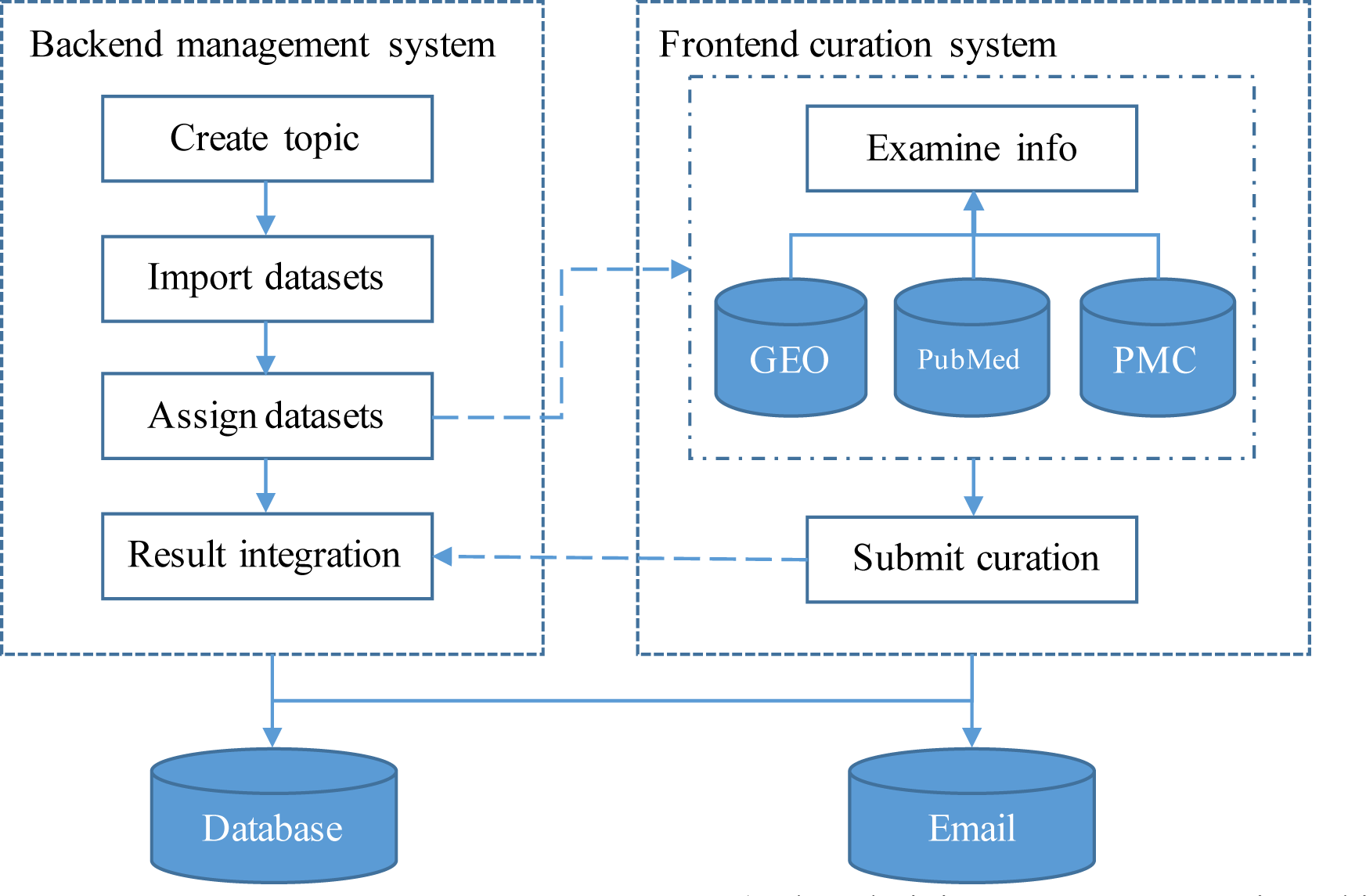
Workflow of GEOMetaCuration. 1) The administrator creates a topic and imports the metadata of a list of datasets to GEOMetaCuration. 2) These datasets are assigned to multiple curators to ensure curation accuracy. 3) The system sends the curators notification emails about the assignments. 4) The curators curate the assigned datasets independently. 5) The curated results are recorded on GEOMetaCuration. 6) The administrator collects the curation results from GEOMetaCuration.

GEOMetaCuration is implemented in Django (https://www.djangoproject.com), a high-level Python web framework. The MySQL database is used for data storage. **Figure 2** shows the database scheme to support multiple curation topics at the same time. To facilitate curating each dataset, a novel design feature combines three types of information on an integrated curation panel for each dataset: the dataset description from GEO, the abstract of the related paper, and the full text. To speed up the recognition of useful information, predefined keywords are highlighted on the panel. GEOMetaCuration is portable and can be customized by other researchers for other curation tasks, such as literature curation. To make it freely accessible, GEOMetaCuration is published under the GNU General Public License V3. Its source code can be freely downloaded at https://bitbucket.com/yubiolab/GEOMetaCuration.

**Figure 2.**
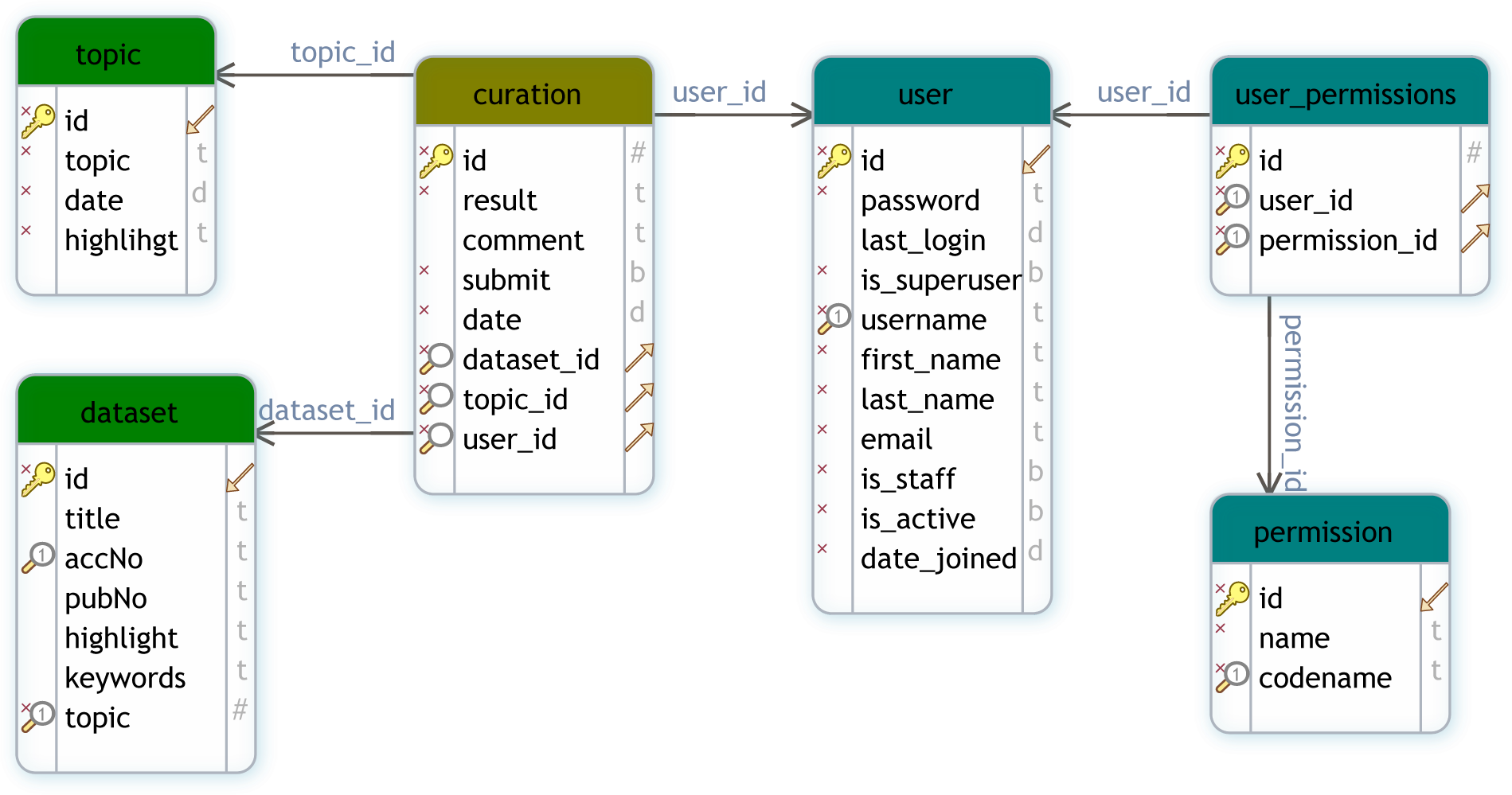
Database schema for GEOMetaCuration. This schema diagram contains six tables that can be classified into three groups: *user management* (user, permission, user_permission), *dataset* (dataset and topic), and *curation,* which contains the curation result of each dataset by each curator.

## Results

### Features and interface of GEOMetaCuration

To facilitate manual curation of GEO metadata, GEOMetaCuration has a feature to automatically load and cache related webpages, including GEO dataset description webpages, PubMed abstract webpages, and PubMed Central (PMC) full text webpages. **Figure 3** shows the preloaded webpages for the GEO dataset GSE10192. The default page of a dataset is the GEO description page, which includes general information such as the experimental design, number of samples, and measurement platform. For example, **Figure 3a** shows the GEO dataset description page for GSE10192, a mouse microarray dataset with the treatment of rosiglitazone. **Figure 3b** shows the preloaded PubMed abstract webpage for the corresponding reference of this dataset, which can allow curators to inspect more-detailed information. **Figure 3c** shows the corresponding full text webpage from PMC, delivering the complete experimental information for ultimate curation decisions. GEOMetaCuration also automatically caches the webpages to save time spent repeatedly requesting the same GEO, PubMed, and PMC webpages. The preloading and caching features can expedite the curation process.

**Figure 3.**
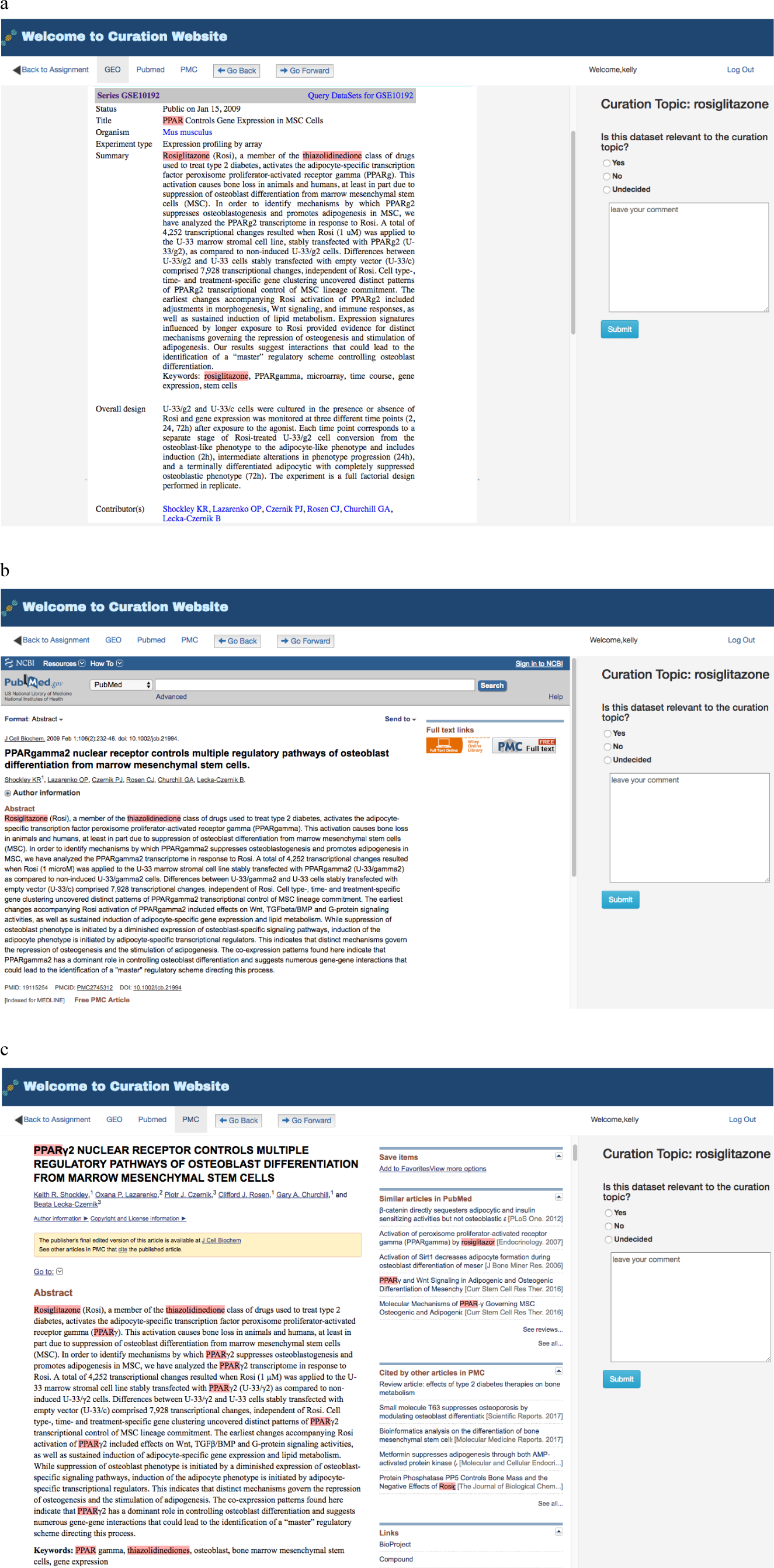
Preloaded GEOMetaCuration webpages for the metadata curation of GSE10192. To expedite metadata curation, GEOMetaCuration is designed to preload webpages for curators to avoid waiting for these webpages to open. To assist in quickly identifying relevant information, predefined keywords are highlighted on the webpages. For the example of GSE10192, keywords are highlighted on three preloaded webpages: (a) the GEO dataset description webpage, (b) the PubMed abstract webpage of the corresponding publication, and (c) the PMC webpage for the full paper.

To speed up the recognition of key information needed for curation, GEOMetaCuration includes a feature to highlight predefined keywords automatically. For example, **Figure 3** shows highlighted keywords “rosiglitazone,” “PPAR,” and “thiazolidinedione” for dataset GSE10192. Using these visual cues, curators can look through webpages quickly in order to make curation decisions.

To help manage the complete lifecycle of curation, GEOMetaCuration can group curation tasks into three stages: “New,” “Submitted,” and “Undecided,” shown as three tabs in **Figure 4**. The “New” tab lists the datasets to be curated. The “Submitted” tab lists the datasets that have been curated with a confident decision from the curator. The “Undecided” tab lists the datasets on which the curator has started but has not reached a final decision (usually because the corresponding metadata are complicated and need more detailed examination). These tabs facilitate efficient organization of datasets assigned to curators.

**Figure 4.**
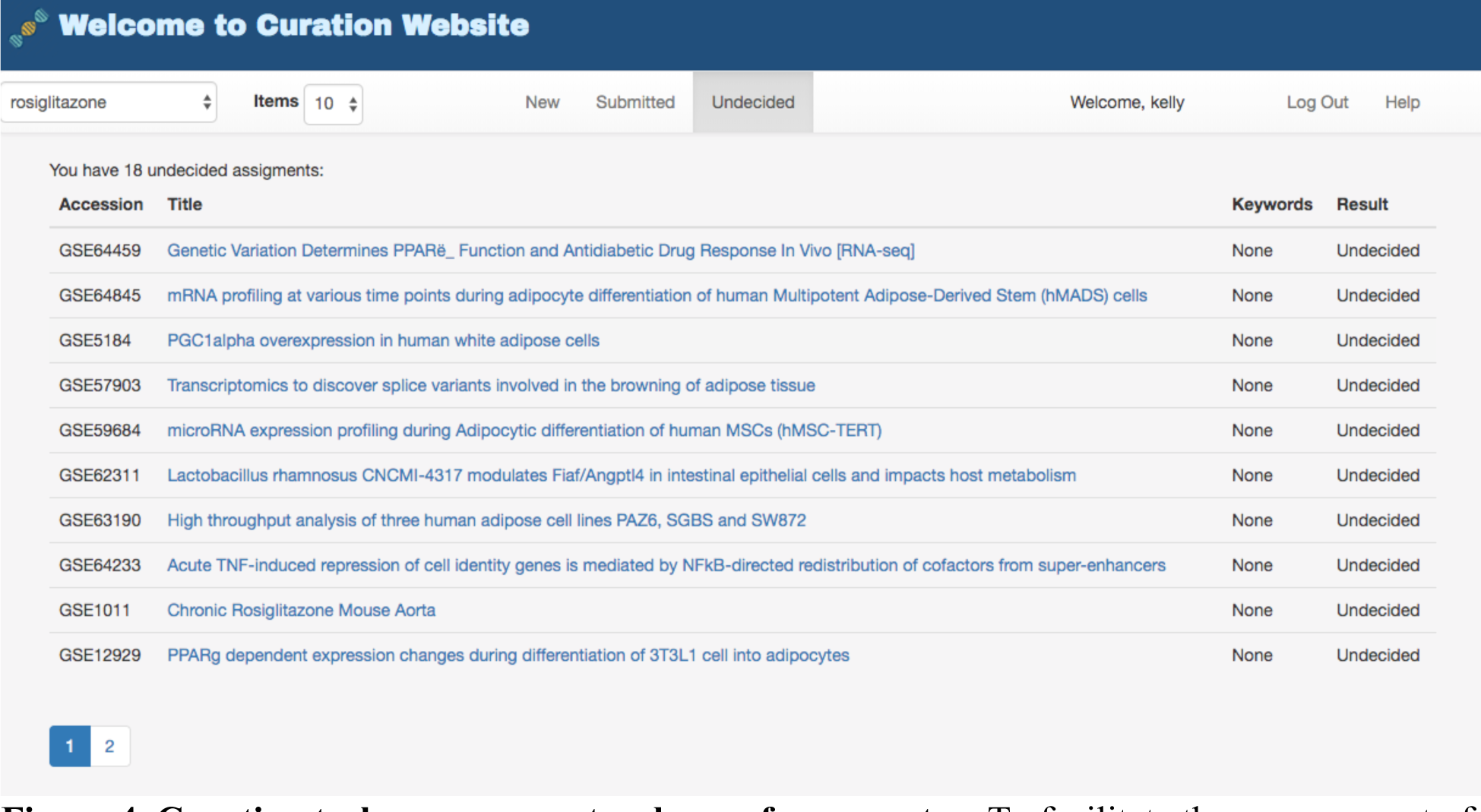
Curation task management webpage for a curator. To facilitate the management of datasets for curation, GEOMetaCuration has three tabs for assigned datasets. The “New” tab shows datasets to be curated. The “Submitted” tab shows datasets that have been curated. The “Undecided” tab shows datasets whose curation results have not been determined yet. Curation topics can be selected on the drop-down menu at the top-left corner (“rosiglitazone” is currently selected in this example). The hyperlink of each dataset links to the corresponding detailed curation webpage, such as the one shown in Figure 3.

To manage the curation task efficiently, GEOMetaCuration fulfills an administration service for different curation tasks. For example, **Figure 5** shows assignment of the curation task for rosiglitazone. The curators and their assigned datasets can be easily selected on the administration webpage. Datasets can be easily assigned to multiple curators at once. This administration service enables powerful management of many curation tasks.

**Figure 5.**
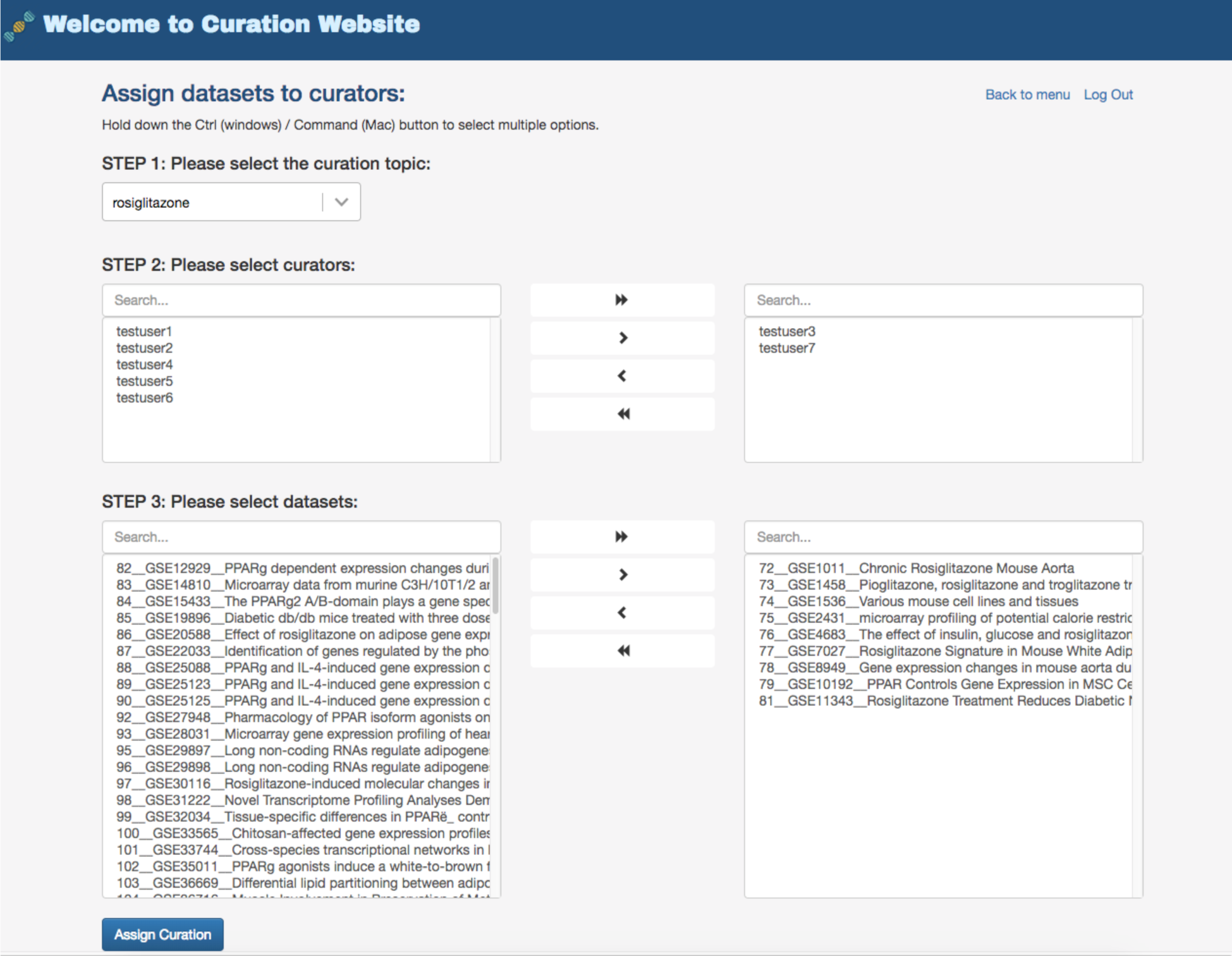
Administration webpage for assigning datasets to multiple curators. The administration webpage allows efficient assignment of datasets to multiple curators for any given specific curation task. Once s/he selects a curation topic, the administrator can then concurrently select the curators and the datasets for an assignment.

### Improved curation time with GEOMetaCuration

We designed the following experiment to confirm improvement to the curation process using GEOMetaCuration. A typical curation process includes two components: 1) performing mechanical operations such as opening the webpages related to each dataset; and 2) looking through the webpages and make a curation decision. Because our tool is designed to only improve the time of the first component and because the time of the second component varies for different curation tasks and for different curators, only the time of the first component was measured and compared.

Google Drive was used in the baseline curation process. Time was recorded for the following activities: 1) copying GEO dataset accession numbers from Google Drive (which contains the GEO gene expression datasets about “rosiglitazone”); 2) Googling the dataset accession numbers and clicking the returned hyperlinks to open the GEO dataset description webpages; 3) clicking the PubMed URLs on these description webpages; 4) clicking the PMC manuscript URLs on the PubMed webpages; 5) typing Ctrl-F and searching for keywords (rosiglitazone, PPAR, and thiazolidinedione) in each webpage opened in steps 2 to 4; and 6) switching back to Google Drive and putting mock results into the rows corresponding to the curated datasets.

Using our curation website, time was recorded for clicking dataset webpages and typing mock results. The time of both with and without the caching feature was recorded to confirm the improvement experience using this feature. **Figure 6** shows the time for each of the first 20 gene expression datasets in **Table S2**. It demonstrates that GEOMetaCuration reduces the time for curation by an average of 60 seconds each time when the information of a dataset is viewed, and the caching feature further reduces it. By reducing this time, the chance for curator distraction is decreased, which can also improve curation efficiency.

**Figure 6.**
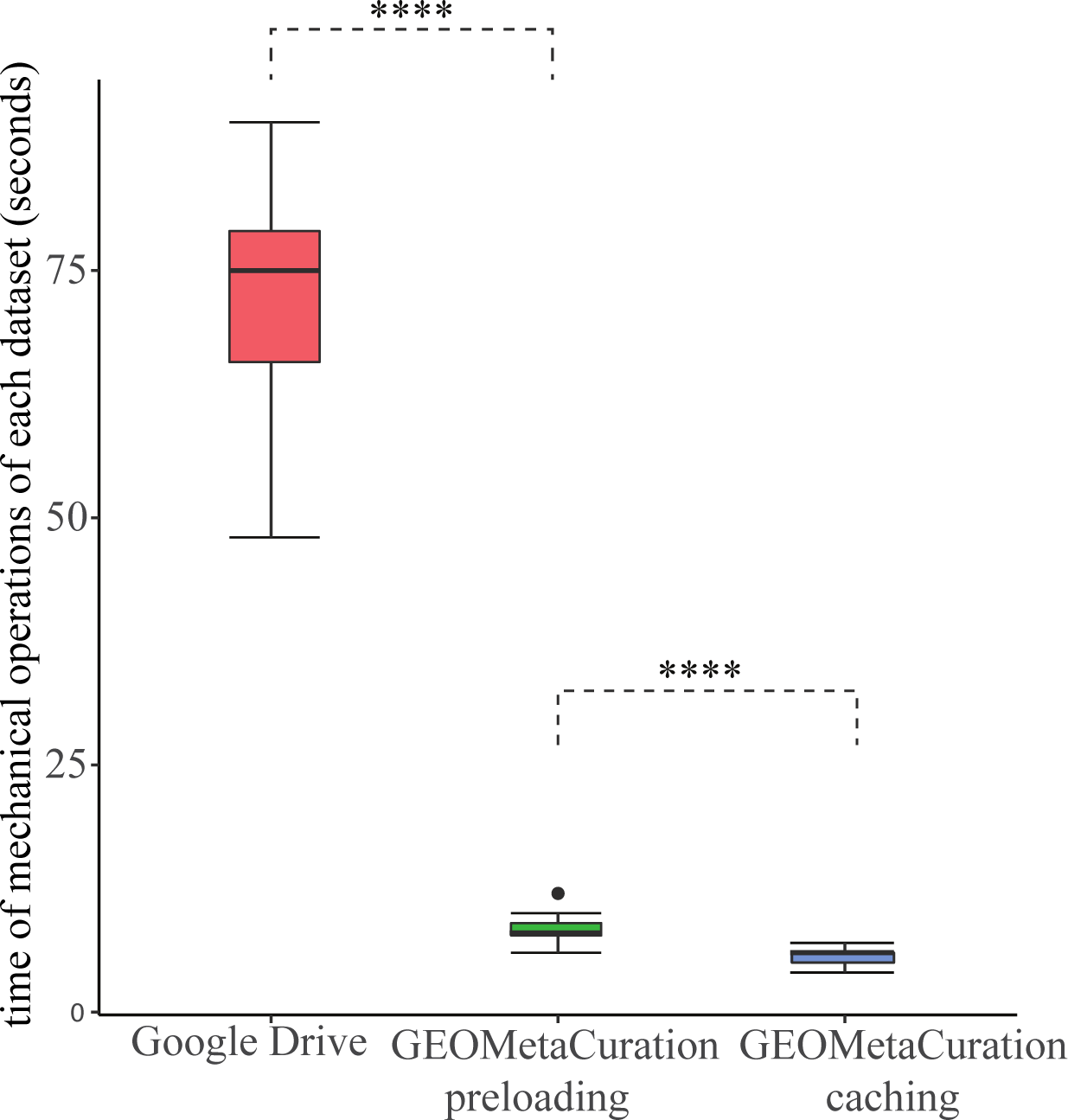
Time spent on mechanical operations required for curation of each of 20 datasets. GEOMetaCuration significantly reduces curation time by eliminating mechanical operations like opening webpages, searching important information, and typing in results. The webpage caching feature further reduces curation time by avoiding repetitive requests to GEO, PubMed, and PMC. **** indicates *p* < 0.0001 (one-sided *t*-test).

### Use case—the curation of human gene expression datasets with rosiglitazone as a treatment

To show the features of GEOMetaCuration, we created a demo website (http://geometacuration.yubiolab.org) using the curation of datasets related to rosiglitazone. Rosiglitazone enhances thermogenesis in brown adipose tissues, which can be helpful to fight obesity and reduce the chances of developing type 2 diabetes (10,11). To study its function in detail, it is critical to collect the complete gene expression data with rosiglitazone as a treatment. The metadata of datasets mentioning “rosiglitazone” were retrieved by querying the database GEO DataSets (https://www.ncbi.nlm.nih.gov/gds) via E-utilities (9) using the following term: (rosiglitazone[Description] OR rosiglitazone[Title]) AND (“Homo sapiens”[porgn: txid9606]) AND gse[Entry Type] AND ((“expression profiling by array”[DataSet Type] OR “non coding rna profiling by array”[DataSet Type]) OR (“expression profiling by high throughput sequencing”[DataSet Type] OR “non coding rna profiling by high throughput sequencing”[DataSet Type])) NOT SuperSeries[All Fields]

The retrieved metadata were first loaded into the MySQL database on the demo website. Then, these datasets were assigned to each curator by the website administrator (**Figure 4**). A notification email was then sent to each curator about the assignment. Each dataset was assigned to multiple curators, and the administrator compared inconsistent answers. To help the curator rapidly locate important information about rosiglitazone, the keywords “rosiglitazone,” “PPAR,” and “thiazolidinedione” were loaded into the back-end management system and were highlighted with different colors on the front end. By examining the metadata of a large number of datasets (Table S2), the final curation results were collected (**Table S1**). This use case confirms the utility of GEOMetaCuration.

## Discussion

Accurate metadata curation is imperative for integrating large-scale biological data to generate biological insights in biomedical research. Since the data for a specific biomedical research problem are usually limited, missing known datasets may result in unnecessary duplication of experiments, leading to increased time and cost and the reduction of research productivity. Meanwhile, although more than 100,000 datasets and more than 2 million samples are hosted on GEO, the number of datasets for a certain biological problem is usually still limited. For example, at the time of writing, a query of general keywords such as “diabetes,” “heart disease,” and “Alzheimer’s disease” in GEO retrieved ~400, ~400, and ~100 human gene expression datasets, respectively. For this limited number of datasets, it would be feasible to manually curate the metadata in just a couple of weeks by well-trained curators. For this reason, we implemented GEOMetaCuration to facilitate the curation process.

Conventional metadata curation is prolonged and inefficient. For example, to construct a metadata database such as RNASeqMetaDB and SFMetaDB, the metadata must be split into multiple files and shared with different curators. Curators then need to manually check the GEO webpages and associated publications. Upon finishing the curation tasks, they submit their intermediate results, which are collected and integrated to construct the final results. It is burdensome to make their notations uniform since different curators have personal habits for marking answers and taking notes. To address these problems, we designed various critical features for metadata curation in GEOMetaCuration. For example, the webpage preloading feature saves curation time by avoiding manually opening webpages related to curation. The keyword highlight feature expedites the identification of relevant information in metadata. Furthermore, the administration service streamlines the management of curators and their tasks.

GEOMetaCuration can enable productive metadata curation of GEO datasets. As its name suggests, GEOMetaCuration currently only supports GEO metadata. But because it is based on the industry-tested web development framework of Django/Python, GEOMetaCuration can be customized for other curation applications, such as literature curation. Its code is open-sourced and can be modified by other researchers for their curation tasks. Therefore, GEOMetaCuration is a useful biocuration application that will eventually contribute to the generation of biological insights using large-scale biological datasets.

## Declarations

### Ethics approval and consent to participate

Not applicable.

### Consent to publish

Not applicable.

## Availability of data and materials

The source code along with its documentation is available on https://bitbucket.com/yubiolab/GEOMetaCuration.

## Authors’ Contributions

PY conceived the general project and supervised it. ZL and PY were the principal developers. ZL, JL, and PY wrote the manuscript. All the authors contributed with ideas, tested GEOMetaCuration, read the final manuscript, and approved it.

## Acknowledgements

The authors thank Bing Jiang for her contribution to GEOMetaCuration.

## References

1. Dauga, D. (2015) Biocuration: A New Challenge for the Tunicate Community. Genesis, 53, 132–142.

2. Cahan, P., Li, H., Morris, S.A., da Rocha, E.L., Daley, G.Q., Collins, J.J. (2014) CellNet: Network Biology Applied to Stem Cell Engineering. Cell, 158, 903–915.

3. Howe, D., Costanzo, M., Fey, P., et al. (2008) Big data: The future of biocuration. Nature, 455, 47–50.

4. Barrett, T., Wilhite, S.E., Ledoux, P., et al. (2013) NCBI GEO: archive for functional genomics data sets-update. Nucleic Acids Research, 41, D991–D995.

5. Bernstein, M.N., Doan, A., Dewey, C.N. (2017) MetaSRA: normalized human sample-specific metadata for the Sequence Read Archive. Bioinformatics, 33, 2914–2923.

6. Kilicoglu, H. (2017) Biomedical text mining for research rigor and integrity: tasks, challenges, directions. Brief Bioinform.

7. Guo, Z.Y., Tzvetkova, B., Bassik, J.M., et al. (2015) RNASeqMetaDB: a database and web server for navigating metadata of publicly available mouse RNA-Seq datasets. Bioinformatics, 31, 4038–4040.

8. Li, J., Tseng, C.S., Federico, A., et al. (2017) SFMetaDB: a comprehensive annotation of mouse RNA splicing factor RNA-Seq datasets. Database-the Journal of Biological Databases and Curation.

9. Agarwala, R., Barrett, T., Beck, J., et al. (2016) Database resources of the National Center for Biotechnology Information. Nucleic Acids Research, 44, D7–D19.

10. Li, Y.L., Li, X., Jiang, T.T., et al. (2017) An Additive Effect of Promoting Thermogenic Gene Expression in Mice Adipose-Derived Stromal Vascular Cells by Combination of Rosiglitazone and CL316,243. International Journal of Molecular Sciences, 18.

11. Ohno, H., Shinoda, K., Spiegelman, B.M., Kajimura, S. (2012) PPAR gamma agonists Induce a White-to-Brown Fat Conversion through Stabilization of PRDM16 Protein. Cell Metabolism, 15, 395–404.

